# Heterogeneity of midgut cells and their differential responses to blood meal ingestion by the mosquito, *Aedes aegypti*

**DOI:** 10.1101/2020.08.31.275941

**Authors:** Yingjun Cui, Alexander W.E. Franz

## Abstract

Mosquitoes are the most notorious hematophagous insects and due to their blood feeding behavior and genetic compatibility, numerous mosquito species are highly efficient vectors for certain human pathogenic parasites and viruses. The mosquito midgut is the principal organ of blood meal digestion and nutrient absorption. It is also the initial site of infection with blood meal acquired parasites and viruses. We conducted an analysis based on single-nucleus RNA sequencing (snRNA-Seq) to assess the cellular diversity of the midgut and how individual cells respond to blood meal ingestion to facilitate its digestion. Our study revealed the presence of 20 distinguishable cell-type clusters in the female midgut of *Aedes aegypti*. The identified cell types included intestinal stem cell (ISC), enteroblasts (EB), differentiating EB (dEB), enteroendocrine cells (EE), enterocytes (EC), EC-like cells, cardia cells, and visceral muscle (VM) cells. Blood meal ingestion dramatically changed the overall midgut cell type composition, profoundly increasing the proportions of ISC and three EC/EC like clusters. In addition, transcriptional profiles of all cell types were strongly affected while genes involved in various metabolic processes were significantly upregulated. Our study provides a basis for further physiological and molecular studies on blood digestion, nutrient absorption, and cellular homeostasis in the mosquito midgut.

## Introduction

Hematophagy is a common trait exhibited by more than 14,000 insect species across five orders, with more than 75% of the known blood-feeders belonging to the Diptera (Adams, 1999). The yellow fever mosquito, *Aedes aegypti* is a well-studied hematophagous dipteran, which transmits human-pathogenic arthropod-borne viruses (arboviruses) including dengue, yellow fever, chikungunya, and Zika viruses thereby posing a major public health threat (Nene et al., 2007). In hematophagous insects, the midgut is the site of blood meal digestion and the organ initiating blood feeding related events, such as vitellogenesis, oogenesis, and also pathogen transmission (Billingsley, 1990). Midgut cells mount innate immune defenses against pathogenic microorganisms, actively shape the gut microbiome, and produce signaling molecules to regulate their own physiology as well as that of other mosquito organs (Caccia et al., 2019). The mosquito midgut is the initial organ that gets infected with an arbovirus. Midgut infection and escape barriers are two important barriers to systemic arbovirus infection determining vector competence of a mosquito for a virus (Franz et al., 2015). The mosquito midgut is an elongated tube-like organ that can be divided into two major regions based on function and anatomy: the anterior midgut and the posterior midgut (Billingsley, 1990). The anterior midgut is primarily responsible for carbohydrate digestion, for example, sugar containing meals, which initially are stored in the mosquito’s crop. The posterior midgut is specialized in blood meal digestion. The midgut organ consists of a single-layered epithelium, which is surrounded by muscle cells, tracheoles and fibroblasts embedded in an extracellular matrix containing banded collagen fibrils (Billingsley and Lehane, 1996).

To obtain a better understanding and overview about midgut cell composition, differentiation, and function in the mosquito it is helpful to first have a look at the situation in the model dipteran, *Drosophila*. The *Drosophila* midgut epithelium is completely renewed every 1–2 weeks with the help of differentiating intestinal stem cells (ISC), which are situated in the midgut (Chen et al., 2016). The adult midgut epithelium of *Drosophila* is composed of four different cell types: intestinal stem cells (ISC), undifferentiated progenitor cells designated enteroblasts (EB), specialized absorptive enterocytes (EC), and secretory enteroendocrine cells (EE) (Nászai et al., 2015). The EC is the predominant cell type and is responsible for digestive enzyme production and nutrient absorption. EE are chemosensory cells that perform important regulatory activities in response to food intake, nutrients, and metabolites through the production and secretion of neuropeptides/peptide hormones. ISC permanently divide and differentiate into EC or EE. ISCs can undergo symmetric and asymmetric divisions to give rise to either two new stem cells or two differentiated daughter cells via the undifferentiated progenitor EB, thereby maintaining homeostasis of the midgut tissue during growth, development, or injury (Nászai et al., 2015, Caccia et al., 2019). Previously, two studies reported that the compartmentalization of the *Drosophila* midgut is associated with distinct morphological, physiological, histological and genetic properties (Buchon et al., 2013, Marianes and Spradling, 2013). Single-cell RNA sequencing (scRNA-Seq) has recently emerged as a powerful tool to monitor global gene regulation in thousands of individual cells allowing the discovery of new cell types and their individual physiological conditions, and to trace their developmental origins (Trapnell, 2015). Guo and colleagues (2019) used scRNA-Seq to identify 10 major EE subtypes in the *Drosophila* midgut producing around 14 different classes of peptide hormones in total while co-producing 2–5 different classes of peptide hormones on average. Furthermore, the authors discovered that transcription factors such as Mirr and Ptx1 were defining regional EE identities resulting in the specification of two major EE subclasses. Hung and colleagues (2020) then presented the first cell atlas of the adult *Drosophila* midgut using scRNA-Seq. It contains 22 distinct cell clusters representing ISCs, EBs, EEs, and ECs. Gene expression signatures of these different cell types were identified and specific marker genes assigned.

In the basal epithelium of the *Drosophila* midgut, ISCs express the ligand *Delta*, which activates the *Notch* signaling pathway in the daughter cells (Ohlstein and Spradling, 2006, Guo and Ohlstein, 2015). *Notch* is a membrane-bound transcription factor regulating stem cell maintenance, cell differentiation, and cellular homeostasis. Specifically, *Notch* signaling is controlling the balance between self-renewing stem cells and their differentiating progeny and determining the type of progeny (Ohlstein and Spradling, 2007). A daughter cell exhibiting a high level of *Notch* activity in the *Drosophila* midgut develops into an intermediate (EB), which then further differentiates into an EC (Perdigoto et al., 2011, Biteau and Jasper, 2014). Low *Notch* expression in combination with high *Delta* and *Prospero* expression levels causes the daughter cell to differentiate into a pre-EE and then further into a mature EE (Zeng and Hou, 2015). *Prospero* is a transcription factor promoting EE specification and therefore qualifies as an EE specific marker. The WT-1 like transcription factor *Klumpfuss* (*Klu*) transiently controls the fate of EB following *Notch* activation, restricting EB to develop into EC but not into EE (Korzelius et al., 2019). Consequently, a loss of *Klu* function results in differentiation of EBs into EEs. Recently, *Klu* was identified in *Drosophila* as a novel marker of midgut-associated EBs (Hung et al., 2020). The *nubbin (nub)/POU domain protein 1* (*Pdm1*) gene is a member of the class II POU transcription factor family (Holland et al., 2007, Tantin, 2013), which is strongly expressed in midgut EC of *Drosophila* to maintain cellular homeostasis. *Nubbin/Pmd1* has been identified as a marker for EC (Hung et al., 2020). Neuropeptides/peptide hormones are essential for the regulation of behavioral actions associated with feeding, courtship, sleep, learning and memory, stress, addiction, and social interactions (Schoofs et al., 2017). In *Drosophila*, midgut EE produce different peptide hormones including *Tachykinins* (*Tk*), *neuropeptide F, (NPF), CCHamide-1* and -*2, allatostatins A* and *C, diuretic hormone 31*, and others, which act on complex neurological circuits affecting locomotion and food search, gut motility, nociception, aggression, metabolic stress, and sleep-feeding regulation (Veenstra et al., 2008, Chung et al., 2017, Hung et al., 2020). Previously, similar neuropeptides were identified in the midgut of *Ae. aegypti* via mass spectrometric profiling (Predel et al., 2010). Apparently, the neuropeptides were found to be unevenly distributed along the midgut. For example, the authors observed that the *short neuropeptide F* (*sNPF*) was most abundantly present in the anterior portion of the midgut.

Our idea was to analyze the gene expression profiles of midgut cells of *Ae. aegypti* females at the single-cell level in order to identify the different cell types present in the female midgut and to reveal how these various cell types respond to the presence of a blood meal. When preparing tissue samples for scRNA-Seq, it has been shown that cell dissociation efficiency and cell viability are factors strongly affecting the overall quality and precision of the analysis (van den Brink et al., 2017). To avoid the problem, transcriptional profiles in single nuclei rather than cell cytoplasm can be analyzed without affecting the overall sensitivity and resolution of the analysis. In particular, this becomes a helpful alternative when processing very minuscule fragile tissues or frozen tissue material (Ding et al., 2020). Accordingly, we used 10x Genomics based single-nucleus RNA sequencing (snRNA-Seq) to generate an atlas of the midgut cell composition in female *Ae. aegypti* and analyzed individual midgut cell expression patterns in response to a blood meal as opposed to a sugar meal.

Here we provide a thorough data analysis demonstrating the overall quality of our snRNA-Seq experiment. We reveal the cell-type composition in the midgut of *Ae. aegypti* and assign marker genes and functions to the various cell types. Our analysis shows possible interactions between midgut cell types and explains the different stages during cell type development. We show that bloodmeal ingestion by a female mosquito dramatically affects her midgut’s overall cell type composition and cell-type specific gene expression patterns.

## Results

### Midgut cell specific snRNA-Seq analysis achieves a > 93 % coverage of the entire *Ae. aegypti* transcriptome

Using 10xChromium technology, our snRNA-seq study was performed on total RNA extracted from single midgut cell nuclei, which were obtained from blood-fed and sugar-fed *Ae. aegypti*. Following selection for high quality nuclei, 4,513 and 5,420 nuclei from the blood-fed and sugar-fed midgut cells, respectively remained for the snRNA-Seq analysis. We obtained 408,155,166 total sequence reads from the midgut cell nuclei of the sugar-fed mosquitoes resulting in an average depth of 65,525 reads per cell, a median of 795 genes per cell and a transcriptome mapping ratio of 83.1 % (**Table S1**). From the midgut cell nuclei of the blood-fed *Ae. aegypti*, 366,541,833 total sequence reads, an average depth of 68,808 reads per cell, a median of 1,226 genes per cell and a transcriptome mapping ratio of 82.3% were recovered. On average, 13,976 and 13,684 protein-encoding genes were detected in individual midgut cell nuclei of the blood-fed and sugar-fed females, respectively. This accounted for 93.6 % to 95.6 % of the annotated genes in the *Ae. aegypti* genome, containing 14,613 protein-encoding genes in total (Matthews et al., 2018). These sequencing data combined with sequencing depth, overall coverage, and transcriptome mapping ratio demonstrate the robustness of our snRNA-Seq experiment providing a reliable basis for the further analysis.

### Unbiased snRNA-Seq analysis identified 20 different cell clusters in the midgut of an *Ae. aegypti* female

The canonical correlation analysis in Seurat (Butler et al., 2018) was used to align the two snRNA-Seq datasets from midguts of blood-fed and sugar-fed females. The integrated snRNA-Seq analysis identified 20 cell clusters visualized with UMAP (Becht et al., 2018) (**Fig. 1, Fig. S1**). These cell clusters have different marker genes **(Table 1, Table S2)**, although there were also common markers identified among some of the clusters. Most of the marker designations were derived from the midgut cells of *Drosophila* and several markers were also derived from other references (Guo et al., 2919, Lin et al., 2008, Dutta et al., 2015, Hung et al., 2020).

**Figure 1.**
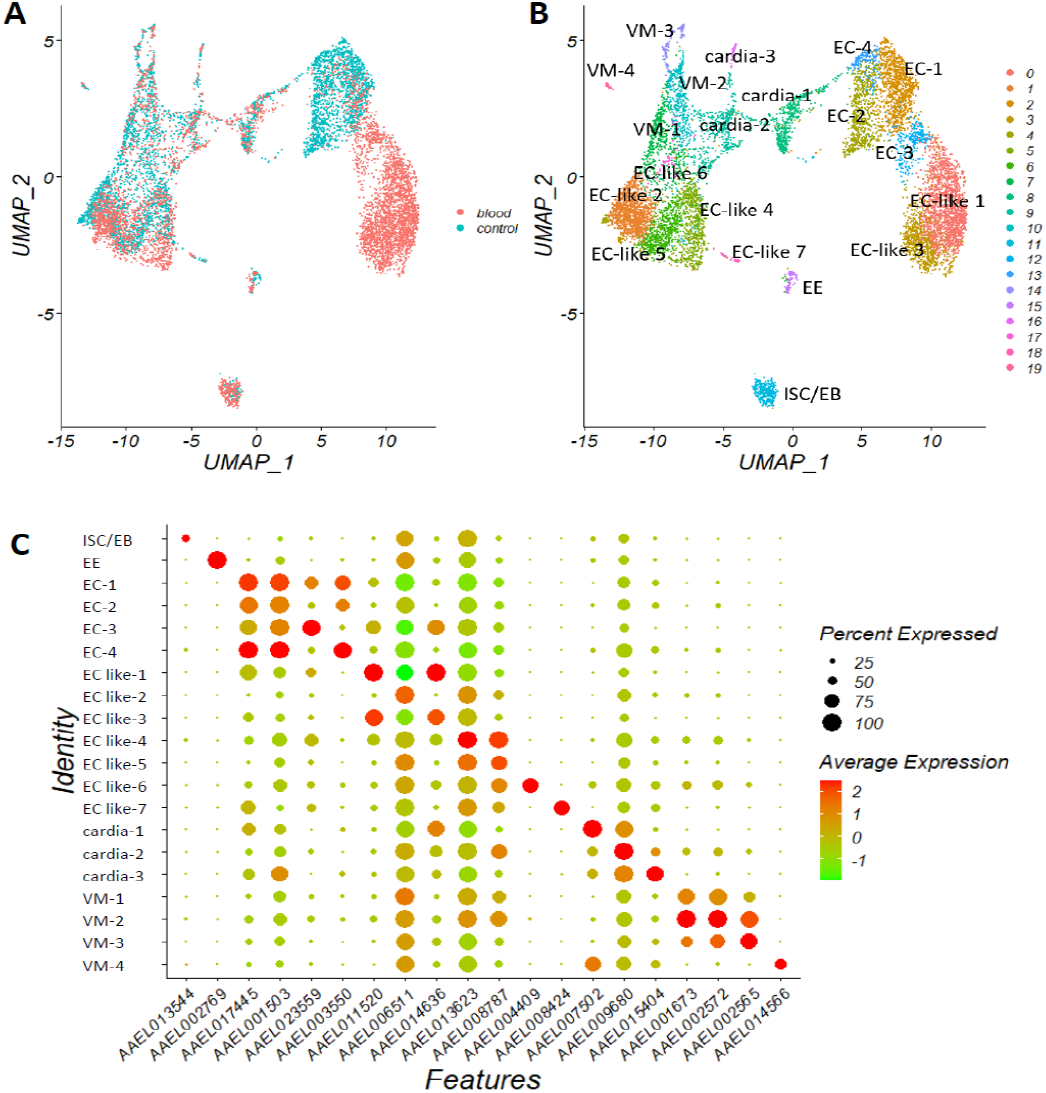
snRNA-Seq identifies 20 cell clusters in the female *Ae. aegypti* midgut. (**A**) UMAPs from midguts of blood-fed (in red) and sugar-fed (in blue) mosquitoes. (**B**) Integrated UMAP based on the datasets obtained from the midguts of blood-fed and sugar-fed mosquitoes showing cell type specific labeling. (**C**) DotPlot showing the proportion of midgut cells expressing marker genes and marker gene expression levels (average lnFoldChange) in each cluster.

**Table 1.** The 20 cell clusters identified in the female midgut of *Aedes aegypti*

| Cluster | cell type | Cell marker | Blood-fed midguts |  | Sugar-fed midguts |  | Fold change | P value |
| --- | --- | --- | --- | --- | --- | --- | --- | --- |
|  |  |  | No. of cells | Proportion | No. of cells | Proportion |  |  |
| cluster_0 | EC like-1 | *Nubbin, sucrose transport protein* | 1691 | 31.2 | 7 | 0.2 | 241.57 | <0.0001 |
| cluster_1 | EC like-2 | *Nubbin, ubiquitin* | 576 | 10.6 | 614 | 13.6 | 0.94 | <0.0001 |
| cluster_2 | EC-1 | *Nubbin,* | 206 | 3.8 | 795 | 17.6 | 0.26 | <0.0001 |
| cluster_3 | EC like-3 | *Nubbin, rhoGTPase* | 861 | 15.9 | 3 | 0.1 | 287.00 | <0.0001 |
| cluster_4 | EC-2 | *Nubbin* | 49 | 0.9 | 737 | 16.3 | 0.07 | <0.0001 |
| cluster_5 | EC like-4 | *Nubbin, trypsin* | 353 | 6.5 | 266 | 5.9 | 1.33 | 0.204 |
| cluster_6 | EC like-5 | *Nubbin, V-type proton ATPase* | 270 | 5.0 | 341 | 7.6 | 0.79 | <0.0001 |
| cluster_7 | VM-1 | *Actin, myosin* | 139 | 2.6 | 400 | 8.9 | 0.35 | <0.0001 |
| cluster_8 | cardia-1 | *Sugar transporter, IRX* | 221 | 4.1 | 276 | 6.1 | 0.80 | <0.0001 |
| cluster_9 | cardia-2 | *Sugar transporter, gambicin* | 227 | 4.2 | 257 | 5.7 | 0.88 | <0.0001 |
| cluster_10 | VM-2 | *Myosin, actin* | 144 | 2.7 | 238 | 5.3 | 0.61 | <0.0001 |
| cluster_11 | ISC/EB | *Delta, Klu* | 227 | 4.2 | 111 | 2.5 | 2.05 | <0.0001 |
| acluster_12 | EC-3 | *Nubbin, angiotensin-converting enzyme* | 278 | 5.1 | 3 | 0.1 | 92.67 | <0.0001 |
| cluster_13 | EC-4 | *Nubbin, aquaporin* | 10 | 0.2 | 152 | 3.4 | 0.07 | <0.0001 |
| cluster_14 | VM-3 | *Titin, muscle lim* | 46 | 0.9 | 89 | 2.0 | 0.52 | <0.0001 |
| cluster_15 | EE | *Prospero* | 49 | 0.9 | 63 | 1.4 | 0.78 | 0.021 |
| cluster_16 | cardia-3 | *C-type lysozyme, gambicin* | 26 | 0.5 | 47 | 1.0 | 0.55 | 0.001 |
| cluster_17 | EC like-6 | *Nubbin, yellow protein* | 20 | 0.4 | 39 | 0.9 | 0.51 | 0.001 |
| cluster_18 | EC like-7 | *Lipase, trypsin* | 15 | 0.3 | 39 | 0.9 | 0.38 | <0.0001 |
| cluster_19 | VM-4 | *Wingless, Dpp, WNT4* | 12 | 0.2 | 36 | 0.8 | 0.33 | <0.0001 |

Note: cardia = cardia cell, EB = enteroblast cell, EE = enteroendocrine cell, EC = enterocyte, EC-like = enterocyte-like cell, ISC = intestinal stem cell, VM = visceral muscle cell

One cluster (# 11), accounting for 2.5 % and 4.2 % of the midgut cells in sugar-fed and blood-fed mosquitoes, respectively, was identified as ISC/EB based on the high expression levels of both, the ISC cell marker gene *Delta* (AAEL025606) and the EB marker gene *Klumpfuss* (*Klu*, AAEL013544) **(Fig. 1, Fig. S2, Table 1, Table S2)**. ISCs and EBs were inseparable in our UMAP. Another cluster (# 15), accounting for 1.4 % and 0.9 % of the midgut cells in sugar-fed and blood-fed mosquitoes, respectively, was designated EE based on high expression of the marker gene *Prospero* (AAEL002769) **(Fig. 1, Fig. S3, Table 1, Table S2)**. Four clusters (# 2, 4, 12, and 13) comprised of EC (EC-1, EC-2, EC-3, and EC-4) based on the strong expression of the marker gene *Nubbin/Pdm1* (AAEL017445) (Pinto et al., 2018) **(Fig. 1, Fig. S4, Table 1, Table S2)**. In seven clusters (# 0, 1, 3, 5, 6, 17, and 18), *Nubbin* expression was clearly lower than in the EC cells of clusters 2, 4, 12, and 13, but still significantly higher than in other cell types. Thus, we designated the cells of clusters 0, 1, 3, 5, 6, 17, and 18 as “EC-like” cells (EC-like 1-7). Cells of clusters 8 (cardia-1), 9 (cardia-2), and 16 (cardia-3) were identified as cardia cells exhibiting significant expression levels of the innate immunity-associated *gambicin* (AAEL004522) and *C-type lysozyme* (AAEL015404) in cardia-2 and cardia-3 clusters or significant expression of two sugar transporters (AAEL010478, AAEL010479) in cardia-1 and cardia-2 clusters (**Fig. 1, Fig. S5, Table 1, Table S2**).

The basal surface of the midgut cells possesses a network of longitudinal and circular muscles that upon contraction can produce peristaltic waves. These contractions serve to move the food along the gut and stir the midgut contents during digestion (Messer and Brown, 1995, Vo et al., 2010). Three clusters (# 7, 10, and 14) were assigned as visceral muscle cells (VM-1, VM2, and VM-3) based on significant gene expression levels for muscle structural proteins, such as thin filament (actin, AAEL001673), thick filament (myosin regulatory light chain 2, AAEL002572), myofilin (AAEL010205), and titin (AAEL002565) (Hartshorne and Gorecka, 2011) **(Fig. 1, Fig. S6, Table 1, Table S2)**. Unlike clusters 7, 10, and 14, cluster 19 (VM-4) did not significantly express muscle protein encoding genes. Instead, *Dpp* (AAEL001876), *WNT4* (AAEL010739) and two *Wingless* protein encoding genes (AAEL014566, AAEL008847) were strongly expressed (**Fig. S6**), which are recognized as marker genes of visceral muscles in the *Drosophila* midgut (Lin et al., 2008, Dutta et al., 2015). We speculate that cells of cluster 19 might represent VM satellite cells (Relaix et al., 2012).

### Gene-expression signatures of the various cell types

The cell type markers described above (average logFC > 0, base = e) were used to analyze the gene-expression signature of each cluster using the GO enrichment analysis program gProfiler (Reimand et al., 2016). Obtained GO terms were then further streamlined using REVIGO (Supek et al., 2011). The cluster ISC/EB in *Ae. aegypti* is similar to its counterpart in *Drosophila* as ISC and EB were not separable in both insects (Hung et al., 2020). Furthermore, gene expression levels of *Delta* and *Klu* were not significantly different between midguts of sugar-fed and blood-fed mosquitoes **(Fig. S2)**. Based on the gene expression profiles of *Delta* and *Klu*, the ISC/EB cluster could be further divided into four sub-groups **(Fig. 2A-D)**. In the first sub-group accounting for 43.2 % of the ISC/EB cells in midguts of sugar-fed mosquitoes, both *Delta* and *Klu* were strongly expressed (**Fig. 2B-D**). In the second sub-group accounting for 48.7 % of the ISC/EB cells in midguts of sugar-fed mosquitoes *Klu* but not *Delta* was highly expressed (**Fig. 2B-D**). In the two other remaining sub-groups accounting for < 10 % of the ISC/EB cells from midguts of sugarfed mosquitoes, both *Delta* and *Klu* were expressed only at very low levels. We therefore identified these two sub-groups as differentiating EB, dEB1 and dEB2. GO enrichment analysis indicated that processes regarding regulation of signaling pathways such as Notch, Insulin, apoptosis, reactive oxygen species, G-protein coupled receptor, lysosomal transport, dephosphorylation, actin cytoskeleton organization, cell adhesion, cell communication, establishment of planar polarity embryonic epithelium, and carbohydrate & lipid metabolism were enriched in the ISC/EB cluster (**Table S3**). These enriched processes, which are known to be essential for maintenance, proliferation, self-renewal and differentiation of the intestinal or other stem cells, indicate that the molecular signature of the *Ae. aegypti* midgut ISC/EB cluster was in accordance with the canonical ISC and EB function in the mosquito midgut.

**Figure 2.**
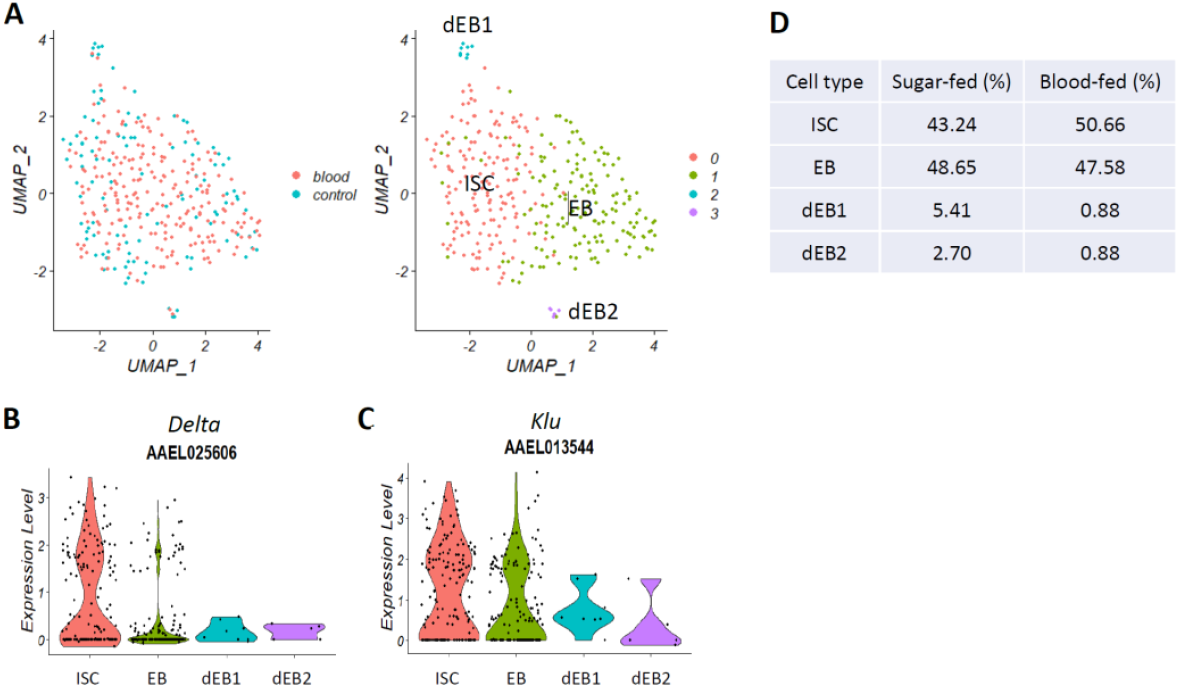
The intestinal stem cell/enteroblast (ISC/EB) cluster in the female *Ae. aegypti* midgut. (**A**) Expression of the markers *Delta* (in red) and *Klu* (in green) within the ISC/EB cluster in midguts from blood-fed and sugar-fed mosquitoes. (**B**) *Delta* and (**C**) *Klu* expression levels in ISC, EB, and differentiating EB (dEB-1, dEB-2) clusters. (**D**) Changes in the overall proportions of ISC, EB, dEB-1, and dEB-2 in the midgut due to blood meal ingestion.

All EEs strongly expressed the marker gene *Prospero* (**Fig. 3A, Fig. S3**). In addition, the neuropeptides/peptide hormones *Tachykinin* (*Tk*), *Neuropeptide F* (*NPF*), *short Neuropeptide F* (*sNPF*), and *CCHamide-1* were specifically expressed in the EE cell cluster (**Fig. 3B**). The majority (58.9 %) of the EEs only produced a single type of peptide hormone/neuropeptide, while around one third (30.3 %) of the EEs co-produced 2-4 peptide hormones/neuropeptides **(Figs. 3B, 3C)**. Interestingly, EEs typically did not co-express *Tk* and *NPF*, and only a very few *NPF* producing EEs co-expressed *sNPF* or *CCHamide-1*. Zeng and colleagues (2015) reported that in *Drosophila*, mature midgut EE cells are generated from a distinct progenitor, preEE, but not from EBs, which develop into EC. In our study, we discovered a few cells in the EE cluster co-expressing *Prospero* and *Delta* **(Fig. 3D)**. However, such *Prospero/Delta* co-expressing cells were not found in the ISC/EB cluster suggesting that the former were in fact preEE cells constituting the progenitor cell type for midgut EE in *Ae. aegypti*.

**Figure 3.**
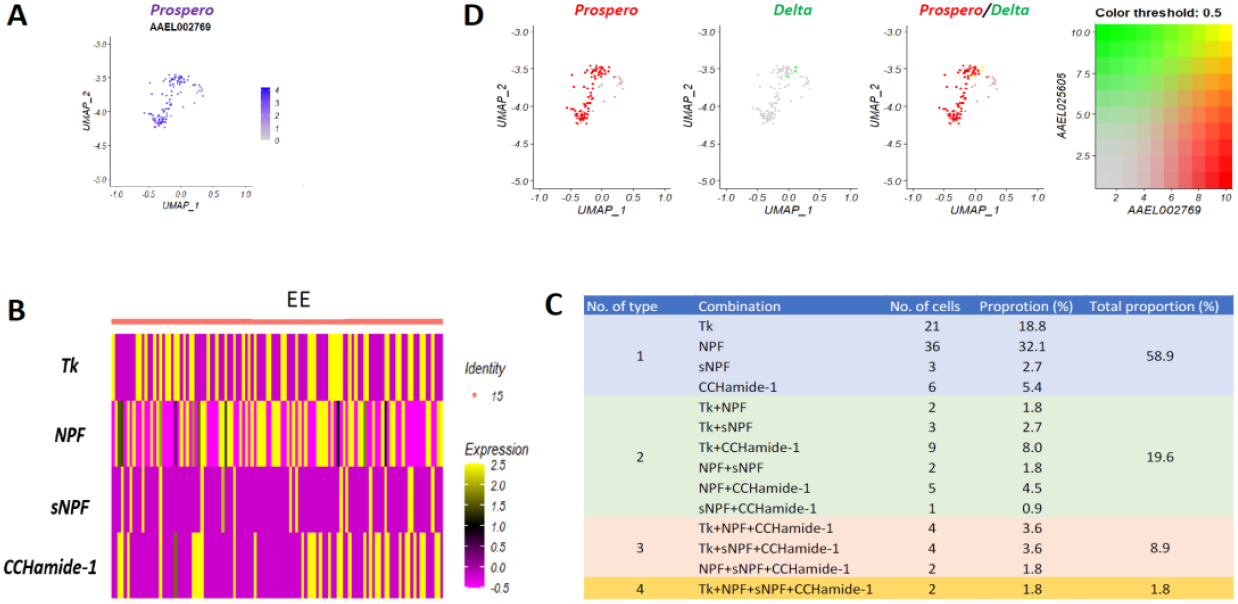
The enteroendocrine (EE) cell cluster in the female midgut of *Ae. aegypti*. (**A**) *Prospero* marker gene expression in the EE cluster as revealed by UMAP. (**B**) Heatmap showing expression patterns of the peptide hormone/neuropeptides *Tk, NPF, sNPF*, and *CCHamide-1* in the EE cluster; expression level = LnFoldChange. (**C**) Proportions of the various peptide hormone/neuropeptide combinations among cell subpopulations within the EE cluster based on their expression levels. *Tk* = *tachykinin, NPF* = *neuropeptide F, sNPF* = *short neuropeptide F*. (**D**) Putative EE progenitor cell subpopulation within the EE cluster expressing the marker genes *Prospero* and *Delta*.

The *Tk*-like receptor, 99D (AAEL006947), was strongly expressed in the cardia-1 cell cluster and in the VM-1, VM-3, and VM-4 clusters (**Fig. 4A**). Within EEs, *Tk* and its receptor were expressed in different cell subpopulations (**Fig. 4B**). The expression of *Tk* and its receptor in different cell types implies that *Tk* functions in the mosquito midgut via a paracrine signaling mechanism. By contrast, the NFP receptor (AAEL010626) was highly expressed within the same EE cell cluster (**Fig. 4C**), albeit in different EE subpopulations (**Fig. 4D**). Expression of the *sNPF receptor* (AAEL013505) was not cell cluster specific **(Fig. 4E)**, however, *sNPF* and *sNPF receptor* expression predominantly occurred in different EE subpopulations **(Fig. 4F).** The *CCHamide-1 receptor* of *Ae. aegypti* has not been identified and annotated so far. EE also express gustatory receptors, which potentially play a role in chemosensation (Park and Kwon, 2011). In *Ae. aegypti*, two gustatory receptors, Gr34 and Gr20, were predominantly expressed in midgut EE where they may be involved in chemosensation together with *neuropeptide FF receptor 2 isoform X2, octopamine receptor*, and *myosuppressin receptor* **(Fig. S7)**, suggesting that midgut EEs are communicating with other cells or tissues. GO enrichment analysis indicated that these chemosensation processes included functions/pathways such as molecule transport, localization, signaling, response to stimulus, cell communication, and G-protein coupled receptor signaling pathway (**Table S4**).

**Figure 4.**
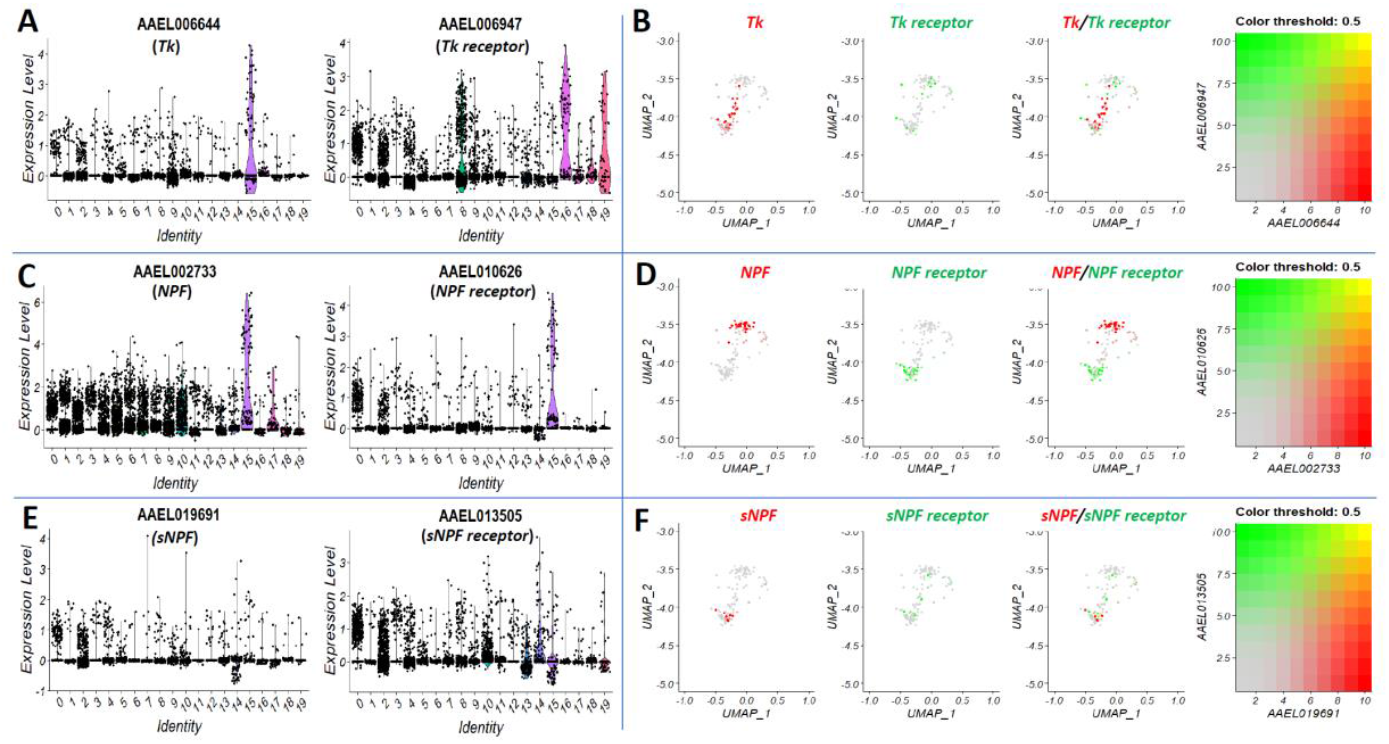
The expression of peptide hormones and their receptors in midgut cells. (**A**) Expression levels of *Tk* (left) and its receptor (right), (**C**) *NPF* (left) and its receptor (right), and (**E**) *sNPF* (left) and its receptor (right) in all cell clusters. (**B**) The expression of *Tk* (in red) and its receptor (in green), (**D**) *NPF* (in red) and its receptor (in green), and (**F**) *sNPF* (in red) and its receptor (in green) within the EE cluster. *Tk* = *tachykinin, NPF* = *neuropeptide F, sNPF* = *short neuropeptide F*.

In all four EC clusters, multiple different metabolic (i.e., protein, macromolecule, nucleic acid, purine ribonucleotide, heterocycle) and biosynthetic (i.e., macromolecule, RNA, nitrogen compound, aromatic compound) processes were enriched (**Table S5**). However, there were differences regarding number and types of the predominant biological processes between these four EC clusters. In five of the seven EC-like clusters (EC like-1, 2, 4, 5, and 7), several of the metabolic and biosynthesis biological processes were also enriched, but not in the other two EC-like clusters (EC like-3, EC-like 6) where responses to stimulus, cell communication, and signaling were predominant instead (**Table S5**). The lipid catabolic process was predominant in cluster 2 (EC-1), whereas the cell component ribosome (40S and 60S ribosomal proteins) was enriched in cluster 1 (EC-like-2). These data indicate that participating in metabolic and biosynthetic processes were the major functions of these EC and EC-like cells, nonetheless, their precise functions seemed to vary.

The cardia, or stomodeal valve, is a specialized fold of the proventriculus that serves as a valve regulating the passage of food material into the anterior midgut and crop (Singh et al., 2011). In the cardia-2 cell cluster, carbohydrate metabolic process was significantly enriched similar to the cardia cluster in *Drosophila* (**Table S6**). The biological processes of response to stimulus, signaling, and cell communication were enriched in the other two cardia cell clusters (cardia-2 and cardia-3) of *Ae. aegypti*. Wnt signaling pathway and actin cytoskeleton organization were only enriched in the cardia-3 cluster (**Fig. S5**).

The cell component sarcomere was highly enriched in clusters VM-1, VM-2, and VM-3 of the four VM clusters **(Table. S2)**. Furthermore, in clusters VM-1 and VM-2, the cell component ribosome was highly enriched. Cells of cluster VM-4 strongly expressed *Dpp, WNT4*, and two wingless genes instead of sarcomere related genes (**Fig. S6**). These molecular signatures are consistent with the proposed functions of the mosquito VM, being responsible for generating peristaltic activity along the underlying midgut epithelium to enable the midgut to ingest a maximal blood meal volume.

### Blood meal ingestion changes the cell composition of the midgut of *Ae. aegypti*

Our snRNA-Seq analysis revealed for the first time that blood meal ingestion induced profound changes on midgut cell type composition affecting 19 of the 20 clusters (the exception was cluster 5, EC-like 4) **(Fig. 1, Table 1).** Due to bloodmeal ingestion, the proportion of the ISC/EB cluster among all midgut cell types increased from 2.5 % to 4 % **(Table 1)**. Likewise, within the ISC/EB cluster, the proportion of ISC increased by ~7 % (from 43.3 % to 50.7 %) whereas the proportions of dEB1 and dEB2 decreased from 5.4 % and 2.7 % to < 1 %. Bloodmeal ingestion did not significantly affect the proportion of EB among midgut cells **(Fig. 2D**).

According to UMAP, the three clusters 0, 3, and 12 (EC like-1, EC like-3 and EC-3) were only apparent in midguts of blood fed mosquitoes but not in those of sugar fed mosquitoes (**Fig. 1**). Accordingly, only three to seven individual cells belonging to these three clusters were detected in midguts of sugar-fed females (**Table 1**). However, 24 h post-blood meal ingestion, their combined proportion increased dramatically from 0.4 % to 52.2 % among all midgut cells. With the exception of cluster 5 (EC-like 4), which was relatively unaffected by blood meal ingestion, the combined proportion of the remaining seven EC and EC-like clusters (EC-1, EC-2, EC-4, EC like-2, EC like-5, EC like-6, and EC like-7) decreased by 39 % (from 60.3 % to 21.2 %) at 24 h post-blood meal ingestion. Thus, in blood fed mosquitoes, there was an overall proportional net gain of EC and EC-like cell clusters among all midgut cells amounting to 13.5 %. Combined with a 1.5 % increase in the proportion of ISC/EB after blood feeding, this implies that at least some of the ECs and EC-like cells must have been generated from EB, whose development from dEB was stimulated by blood meal ingestion (**Table 1**).

The proportion of the EE cluster among midgut cells decreased by 0.5 %, from 1.4 % in sugar-fed females to 0.9 % in blood-fed females. The combined proportion of the three cardia cell clusters (cardia-1, cardia-2, and cardia-3) among all midgut cells decreased by 4% (from 12.8 % to 8.8 %) in midguts of blood-fed females. This is consistent with the current idea that that cardia cells do not play any major role in blood meal digestion. Similarly, the combined proportion of the four VM clusters 7 (VM-1), 10 (VM-2), 14 (VM-3), and 19 (VM-4) decreased by ~10 % from 16.9 % in sugar-fed mosquitoes to 6.3 % in blood-fed mosquitoes (**Table 1**). All these profound bloodmeal induced changes in the midgut cell composition seem to be essential to prepare the organ for efficient blood digestion and increased nutrient absorption.

### Blood meal ingestion dramatically affects the transcriptomes of the various cell types in the midgut of *Ae. aegypti*

In addition to causing profound changes to the proportions of the various midgut cell types, blood meal ingestion also strongly affected the transcriptomes of the cell types (**Fig. 5, Table S7**). Between 26 and 201 protein encoding genes were significantly (fold change >= 2, and p < 0.05) up-regulated in all cell clusters with the exceptions of clusters 0 (EC like-1), 3 (EC like-3), and 12 (EC-3) (**Fig. 5B**). The majority of these up-regulated genes were involved in metabolic processes (organonitrogen compound, carbohydrate and small molecule) and iron ion transport (**Table S8**). In the ISC/EB cluster, additional upregulated genes were involved in biosynthetic processes (organic acid, small molecule and L-serine) and cellular homeostasis. Furthermore, bloodmeal ingestion caused a significant upregulation of *Notch* (AAEL023745) (**Fig. S8**). In EEs, biological processes such as cellular homeostasis and intracellular receptor signaling pathway were significantly enriched in the upregulated genes due to blood meal ingestion (**Table S8**).

**Figure 5.**
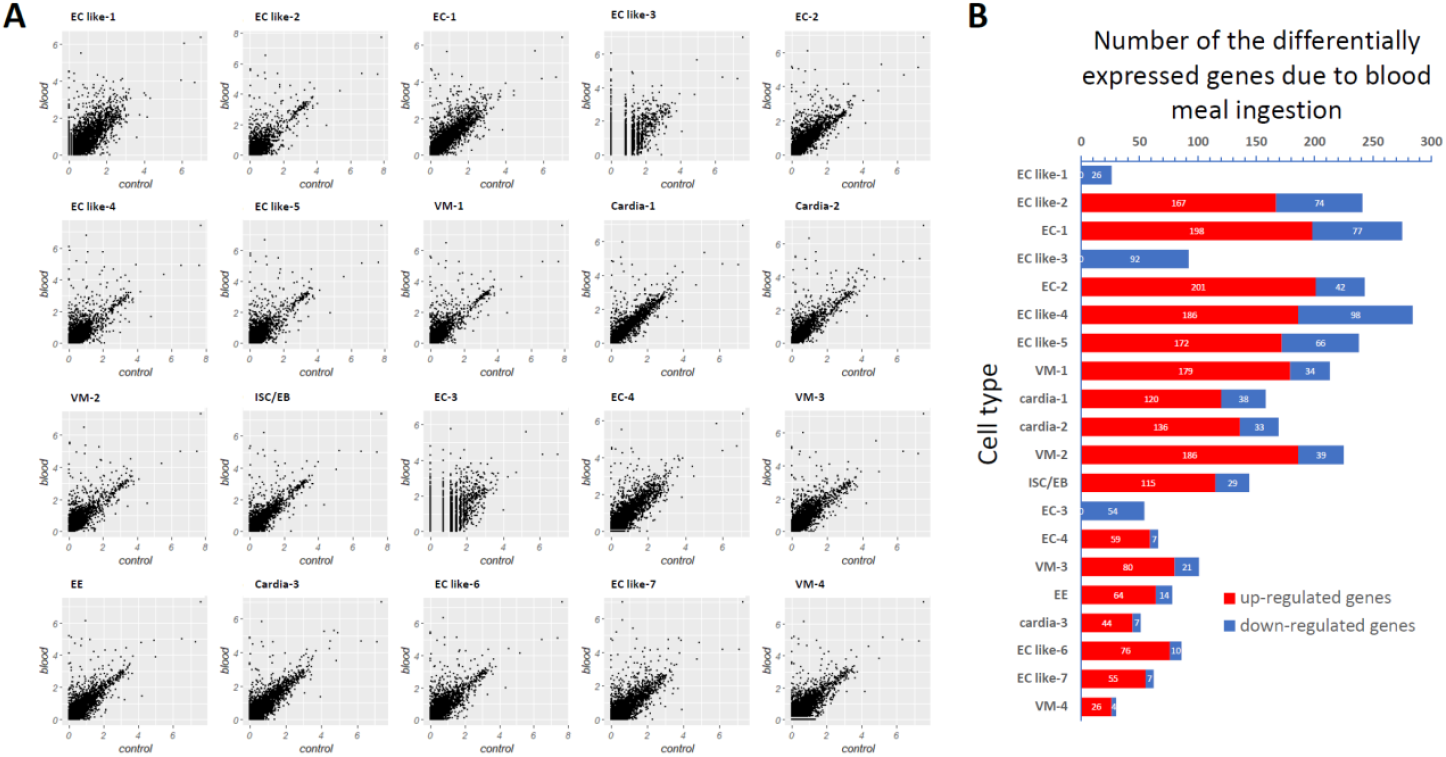
The response of each cell cluster in the midgut of an *Ae. aegypti* female to blood meal ingestion. (**A**) Scatter plot showing the expression levels (average lnFoldChange) of all genes (each gene is represented by a black dot) in each cell cluster of the control group (sugar-fed midgut, x axis) as compared to the blood meal fed group (blood-fed midgut, y axis) at 24 h post-blood meal. Dots on the 45° axis represent genes whose expression levels were unaffected by blood meal ingestion. (**B**) Number of differentially expressed genes in each midgut cell cluster due to blood meal ingestion.

Furthermore, in all EC and EC-like clusters except in clusters EC like-1, EC like-3, and EC-3 biosynthetic processes (organic acid, small molecule and L-serine), oxidation-reduction process, and chemical homeostasis were enriched in addition to metabolic processes and iron ion transport (**Table S8**). Cellular lipid catabolic process was only enriched in the EC-1 cluster. In the cardia cells of all three clusters, carbohydrate metabolic process was significantly enriched during blood meal ingestion. Two biological processes, oxidation-reduction process and cellular response to chemical stimulus, were significantly enriched in two VM clusters (VM-1 and VM-2) of the bloodfed mosquitoes whereas in VM-4 cluster, proteolysis, iron ion transport, and chemical homeostasis were the predominant enriched biological processes. Bloodmeal ingestion increased expression levels of certain genes (i.e., serine-type endopeptidase, serine collagenase, and protein G12) by > 100-fold (**Fig. 6A-C**). Two ferritin subunits were significantly up-regulated as well (> 10-fold) (**Fig. 6D**). Several of these highly up-regulated genes had been earlier described in a whole-midgut RNA-Seq study (Bonizzoni et al., 2011). By comparison, substantially fewer genes (between 4 and 98 genes) were downregulated in midgut cells as a response to blood meal ingestion (**Fig. 5B**). For example, the molecular function hydrolase activity was significantly down-regulated in clusters ISC/EB, VM-3, and VM-4, whereas the biological process proteolysis was downregulated in clusters EE, EC like-6, cardia-3, cardia-4, VM-1, VM-2, and VM-4 (**Table S8**). The cell component ribosome was downregulated in clusters EC-1, EC-2, and EC-3. Overall, trypsin 3A1 (AAEL007818), metalloproteinase M12A (AAEL014516), and aquaporin (AAEL005001) were the most common down-regulated genes in midgut cells in response to blood meal ingestion (**Fig. 6E**).

**Figure 6.**
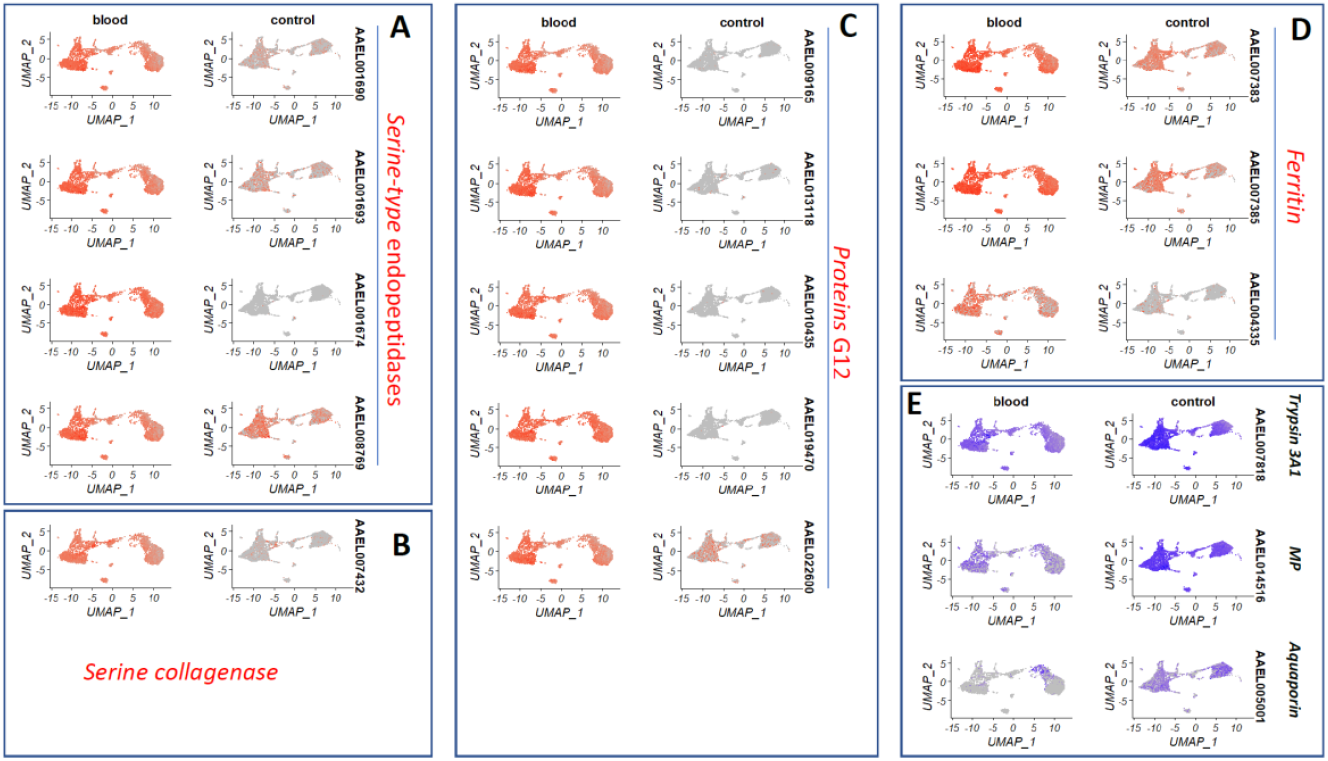
Genes of the female *Ae. aegypti* midgut that responded strongly to blood meal ingestion. (**A**) Four serine-type endopeptidases, (**B**) a serine collagenase, (**C**) five G12 protein encoding genes, (**D**) three ferritins were significantly up-regulated in most of the cell clusters due to blood meal ingestion. (**E**) Trypsin 3A1, M12A seminal metalloproteinase 1 (MP), and aquaporin were significantly downregulated in most of the cell clusters due to blood meal ingestion. Shown color intensities correspond to the relative gene expression levels in the cell nuclei.

## Discussion

The mosquito midgut is essential for nutrition processing and absorption as well as female fecundity due to bloodmeal digestion and corresponding hormone signaling (Sanders et al., 2003). In culicine mosquitoes such as *Ae. aegypti*, the mosquito midgut is also a critical organ determining the vector competence for arboviruses. A wealth of important information regarding the physiology, biochemistry, and molecular biology of the mosquito midgut as-a-whole has been generated over the past decades. For example, previously, it was shown that (blood) meal ingestion causes drastic structural changes to the midgut epithelium of a mosquito (Reinhardt and Hecker, 1973, Houk, 1977, Rudin and Hecker, 1979, Dong et al., 2017, Kantor et al., 2018, Cui et al., 2019). Bloodmeal ingestion also dramatically affects the mosquito midgut’s overall transcriptome and proteome (Sanders et al., 2003; Bonizzoni et al., 2011; Cázares-Raga et al., 2014). However, detailed information on individual midgut cell identity and function has been lacking so far. Here, we revealed for the first time the cellular diversity of the midgut of *Ae. aegypti* using 10xChromium snRNA-Seq and assessed how blood meal ingestion is affecting cell type composition and gene expression patterns in individual midgut cells. Recently, Hung and colleagues performed a scRNA-Seq analysis of the *Drosophila* midgut using the platforms 10x Genomics and inDrop (Hung et al., 2020). 10x Genomics scRNA-Seq detected up to 550 genes per cell (median value) and 9,455 genes in total from 2,723 midgut epithelial cells. By comparison, our 10x Genomics snRNA-Seq analysis enabled us to detect 795 genes per nucleus (median value) and 13,976 genes in total from 6,229 nuclei, which were obtained from the female *Ae. aegypti* midgut. As the entire *Ae. aegypti* genome contains 14,613 protein encoding genes (Matthews et al., 2018), this means that we were able to detect almost all (up to ~96 %) of the annotated genes among the midgut epithelial nuclei/cells. Our study also confirmed that both, scRNA-Seq and snRNA-seq are capable of sufficient gene detection allowing an adequate representation of cell populations (Denisenko et al., 2020, Ding et al., 2020).

Both being dipterans, *Ae. aegypti* and *Drosophila* have numerous traits in common regarding their morphology, physiology, biochemistry, development, and certain behaviors. Single-cell transcriptomics revealed that their midguts share common features as well. For example, both species have similar numbers of distinguishable midgut cell type clusters, 20 clusters for *Ae. aegypti* compared to 22 clusters for *Drosophila*. Furthermore, both species possess a single midgut ISC/EB cluster with common cell type markers, such as *Delta* for ISC and *Klu* for EB. The cell type markers *Prospero* for EE and *Nubbin* for EC were also similar between *Ae. aegypti* and *Drosophila*. Differences regarding midgut cell composition between the two species became obvious when comparing their EE and cardia clusters, which varied in numbers. The three EE clusters found in *Drosophila* as opposed to a single EE cluster in *Ae. aegypti*, produced also different neuropeptides not found in the mosquito. Furthermore, the *Drosophila* midgut contains additional cell types and markers such as *esg* for ISCs, *lab* for middle ECs, *PGRP-SC2* for copper cells/ion cells, *PGRP-SC1a*, and *PGRP-SC1b* for large flat cells, which are absent in the mosquito. Another difference between the two insects lies in the proportion of individual midgut cell types. The *Drosophila* midgut on average consists of 8 % ISCs/EBs, 8 % EEs, and 81 % ECs whereas the midgut of a sugar-fed *Ae. aegypti* female contains 2.5 % ISCs/EBs, 1.4 % EEs, and 66.4 % ECs (in addition to 29 % EC-like cells). The EEs of the *Drosophila* midgut produce up to 15 peptide hormones and ~80 % of individual EEs co-express 2–5 classes of peptide hormones, which exhibit region-specific expression patterns along the length of the midgut (Guo et al., 2019, Hung et al., 2020). This is in contrast to the EEs of *Ae. aegypti*, which generate only four peptide hormones, *Tk, NPF, sNPF*, and *CCHamide-1* with 58.9 % of the EEs producing only a single peptide hormone/neuropeptide while 30.3 % of the EEs co-producing 2-4 peptide hormones/neuropeptides. The dramatic differences between *Drosophila* and *Ae. aegypti* regarding the diversity, quantity, expression levels, and expression pattern of the peptide hormones might be a consequence of the very different food sources both insects are pursuing, which likely are more diverse for *Drosophila*. Earlier, it was reported that in the adult *Drosophila* posterior midgut, EE cells are generated from stem cells through a distinct progenitor (preEE), but not from EBs (Zeng and Hou, 2015). We detected a small number of EEs co-expressing *Prospero* and *Delta* in the *Ae. aeypti* midgut, strongly suggesting that these cells, which we designated as preEEs represent the likely EE progenitor in the mosquito midgut.

We discovered that blood feeding comprehensively changed the overall cell type composition and the transcriptome of each cell cluster. The EC like-1, EC like-3 and EC-3 (clusters 0, 3 and 12) were the most responsive cell-type clusters. Since blood meal digestion is performed in the posterior midgut, which is capable of considerable expansion and is also the major site of vigorous trypsin enzyme activity (Van Handel, 1984, Billingsley and Hecker, 1991), it can be speculated that EC like-1, EC like-3, and EC-3 were located in the posterior midgut. These cells could be mature ECs, which are specialized in blood meal digestion and nutrient absorption. This is also reflected by the drastic increase in their overall proportion in response to a blood meal. However, it seems to be impossible to newly generate these large quantities of ECs and EC-like cells in the midgut just within 24 h pbm. But there is another possibility: instead of being newly generated, these blood meal digesting ECs and EC-like cells could be transformed from other established ECs and EC-like cells in the midgut. This could also explain the dramatic blood meal induced decrease in the proportion of the EC-1, EC-2 and EC-4 clusters, which, based on UMAP, seem to be in close proximity to the EC like-1, EC like-3 and EC-3 clusters. Thus, clusters EC-1, EC-2 and EC-4 may represent immature ECs, which are also situated in the posterior midgut and which can be rapidly transformed into mature blood digesting ECs. Blood meal ingestion also caused an increase in the proportion of the ISC/EB in conjunction with a decrease of the dEBs, suggesting that blood feeding stimulates the cell differentiation of ISC into EB, EB into dEB, and dEB into EC or EC like cells. Thus, blood meal ingestion affected, in a variable manner, almost all of the identified midgut cell clusters.

The majority of the midgut cell genes that were upregulated as a response to blood meal ingestion were involved in the various metabolic processes. These genes included serine-type endopeptidase, serine collagenase, protein G12, and ferritin and had been earlier reported to be blood meal responsive as revealed by several transcriptome analyses on whole midguts (Sanders et al., 2003, Bonizzoni et al., 2011). In our study, midguts of the blood-fed mosquitoes were analyzed at 24 h pbm, a time point of strong digestive enzyme activity and systemic nutrient transport throughout the mosquito organism (Houk and Hardy, 1982). Surprisingly, these genes with metabolic function were also upregulated in ISC/EB and VMs. Future investigations could help to address and explain this phenomenon.

The functional comparison of the midgut cell types between *Drosophila* and *Ae. aegypti* also points to several phenomena that warrant further clarification. For example, in *Drosophila* the intestinal regionalization is defined after adult emergence and it remains stable throughout the remaining life span of the insect (Buchon et al., 2013). The mosquito midgut, by contrast, does not seem to be similarly regionalized as in *Drosophila* (Billingsley, 1990). However, the fact that multiple EC clusters with different expression patterns are present in the *Ae. aegypti* midgut implies the possibility that a certain degree of regionalization may exist in the mosquito midgut, although there is no clear evidence so far. Regardless, our study here lays the foundation for further investigations regarding intestinal regionalization, stem cell regeneration, and cell-type specific pathogen invasion in the midgut of *Ae. aegypti* and other mosquito species.

## Materials & Methods

### Mosquitoes

*Ae. aegypti* mosquito (strain: Higg’s White Eye) larvae were fed on tropical fish food (Tetramin, Melle, Germany). Adult mosquitoes were maintained on raisins and distilled water. Artificial blood feeding was performed using defibrinated sheep blood (Colorado Serum Company, Denver, CO), which was provided in glass feeders covered with Parafilm acting as the feeding membrane. Mosquitoes were reared in an insectary, which is maintained at 28 °C, 80 % relative humidity, and a 12h light/12h dark cycle.

### Isolation of single nuclei from midgut cells

Around 30 midguts were dissected from 7-day old blood-fed female *Ae. aegypti* at 24 hours (h) post-blood meal (pbm) or from sugar-fed females. Dissected midguts were washed in Schneider’s *Drosophila* Medium on ice, followed by centrifugation for 5 min at 500 x *g* and 4 °C. Supernatant was aspirated before the isolation of single nuclei was performed using the Nuclei PURE Prep nuclei isolation kit (Sigma-Aldrich, St. Louis, MO, USA) in conjunction with a modified protocol. Briefly, midguts were resuspended in 100 μl of fresh lysis buffer (Nuclei PURE Lysis Buffer containing 1 mM dithiothreitol (DTT) and 0.1 % Triton X-100), and ground using a micro pestle. Then another 400 μl lysis buffer was added followed by incubation on ice for 15 min. A 900 μl volume of 1.8 M sucrose cushion buffer (Sigma-Aldrich) was mixed with the homogenized midgut tissue. Two new 1.5 ml microcentrifuge tubes containing 500 μl of 1.8 M sucrose cushion buffer were prepared before 700 μl of the solution from the previous step was gently layered on the top of the 1.8 M sucrose cushion. Following centrifugation at 13,000 x *g*, for 45 min at 4 °C, the supernatant was gently aspirated and the pellet resuspended in 500 μl of Nuclei PURE Storage Buffer. Thereafter, the solution was filtered using a cell strainer with a mesh width of 40 μm (Fisher Scientific, Waltham, MA, USA). The filtered solution was then centrifuged at 500 x g, 4 °C for 5 min before the supernatant was aspirated again and the pellet resuspended in 200 μl of Nuclei PURE storage buffer. The survival and quality of the isolated nuclei was checked via the trypan blue staining method. The concentration of the nuclei was determined using a Countess II FL Automated Cell Counter (ThermoFisher, Waltham, MA, USA).

### Conduction of snRNA-Seq using 10x Chromium technology

An estimated number of ~5,000 single nuclei from midguts of the blood-fed or sugar-fed *Ae. aegypti* females was used for scRNA-Seq. Nuclei were processed through the GEM well on a 10x Chromium Controller (10x Genomics, Pleasanton, CA, USA). The two libraries (nuclei from blood-fed midguts versus nuclei from sugar-fed midguts) were then prepared following the user guide and then subjected to 10x Genomics single-cell isolation and RNA sequencing following the manufacturer’s recommendations. A NovaSeq SP PE250 instrument was used for deep sequencing. Library construction and sequencing were performed at the DNA Core Facility of the University of Missouri.

### Analysis of snRNA-Seq data

Cell Ranger (3.1.0) was used to perform the initial data analysis. Briefly, demultiplexing, unique molecular identifier (UMI) collapsing, and alignment to the *Ae. aegypti* transcriptome (AaegL5.2_pre_mRNA) were performed. Raw data generated by Cell Ranger were then imported into the R toolkit Seurat (v3.1) to compare snRNA-seq data sets across different conditions, technologies, or species (Butler et al., 2018). Midgut cell nuclei showing transcription of at least 500 genes and no more than 6500 genes were used to further analysis. A global-scaling normalization method, LogNormalize, was employed to normalize the gene transcription measurements for each cell by its total transcript count, multiplied by the scale factor 10,000 (default) before log-transforming the result. The FindIntegrationAnchors function was used to find anchors between the data of the blood-fed and sugar-fed midguts, these anchors were used to integrate the two datasets together using the IntegrateData function. The default method was used for clustering. Uniform manifold approximation and projection (UMAP) was used to reduce the data dimensionality in order to visualize single-cell data. FindClusters was applied selecting a resolution parameter of 1 for cell clustering. The function FindConservedMarkers was used to identify canonical cell type marker genes that were conserved in the blood-fed and sugar-fed midguts. The DotPlot function including the split.by parameter was used to view conserved cell type markers across conditions, this way showing both the expression level and the percentage of cells in a cluster expressing any given gene. The cell types were identified based on known marker genes of the *Drosophila* midgut (Guo et al., 2019, Hung et al., 2020).

### Analysis of gene expression signatures of each midgut cell type

Using gProfiler, a gene whose expression level in a specific cell-type cluster was higher than its average expression level in all other cell clusters was selected as marker for that specific cell cluster. The same software tool was used to analyze the overall gene expression profiles in the identified cell clusters (Reimand et al., 2016). Obtained GO terms were further analyzed with REVIGO to reduce redundancy, prioritize the enriched/statistically significant terms, and display only their representatives to ease data interpretation (Supek et al., 2011).

### Differential gene expression analysis in response to blood meal ingestion

The function FindMarkers of Seurat was used to identify the differentially expressed genes (fold-change ⩾ 2 or ⩽ 0.5, and adjusted p value < 0.05) between the midguts from sugar-fed and blood-fed mosquitoes. The functions FeaturePlot and VlnPlot of Seurat were used to visualize gene expression changes caused by blood meal ingestion. Significance levels regarding cell proportions in each cluster between midguts from sugar-fed and blood-fed mosquitoes were statistically analyzed via the Wilcoxon signed-rank test.

## Acknowledgements

We would like to thank Drs. Nathan Bivens and Ming-Yi Zhou of the DNA Core of the University of Missouri for their help with the snRNA-Seq library preparation and deep-sequencing. Thanks also to Dr. Christopher Bottoms of the Informatics Research Core of the University of Missouri for pre-processing the raw data and to Jingyi (Jenny) Lin for mosquito rearing. This research was funded by National Institutes of Health - National Institute of Allergy and Infectious Diseases (NIH-NIAID) grant 1R01AI134661. The funding agency was not involved in the design and analysis of this research work.

## Supplemental Tables (Excel format)

**Table S1. Summary of single-nucleus RNA-Seq data.**

**Table S2. The top 10 marker genes for 20 cell clusters of the midgut of *Aedes aegypti*.** The top 10 markers for each cluster are listed, their descriptions, expression levels (LnFoldChange), their first two principle components (pct. 1 and pct.2) and adjusted p values are shown.

**Table S3. Enriched biological processes in the ISC-EB cluster.**

**Table S4. Enriched biological processes in the EE cluster.**

**Table S5. Enriched biological processes in the EC and EC-like clusters.**

**Table S6. Enriched biological processes in the cardia clusters.**

**Table S7. Differential gene expression in each cell cluster as a response to blood meal ingestion.** The differentially expressed genes (fold change ≥ 2 or ≤ 0.5 and adjusted p value < 0.05) of each cluster upon blood meal ingestion are listed. Gene IDs, descriptions, expression levels (LnFoldChange) in the sugar-fed midguts and blood-fed midgut, fold-change expression after blood meal ingestion and adjusted p values are shown.

**Table S8. Enrichment of biological processes in each cell cluster as a response to blood meal ingestion.**

## Supplemental Figures

**Fig. S1.**
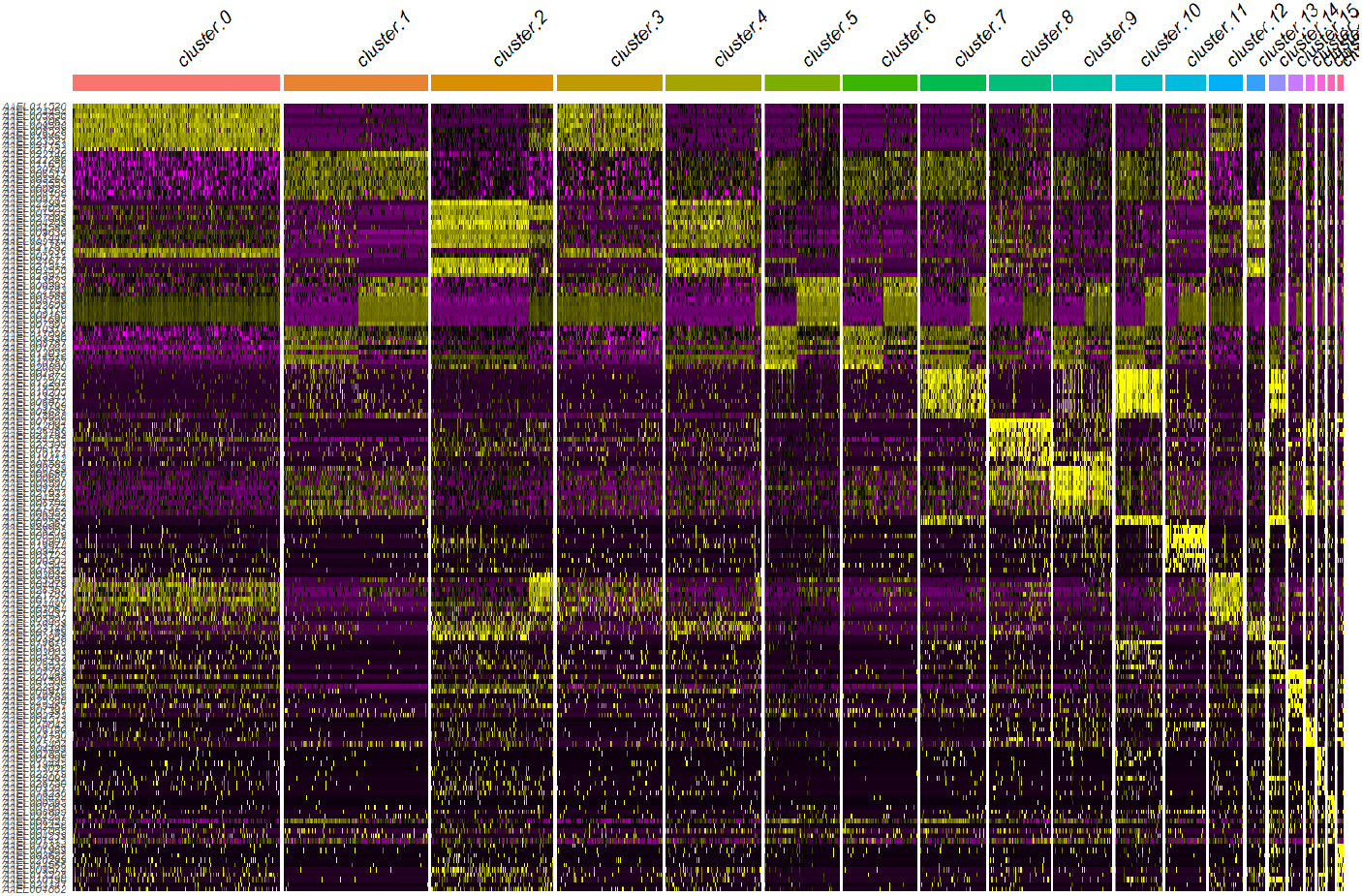
Heatmap showing expression levels (LnFoldChange) of the top 10 marker genes for each cell cluster.

**Fig. S2.**
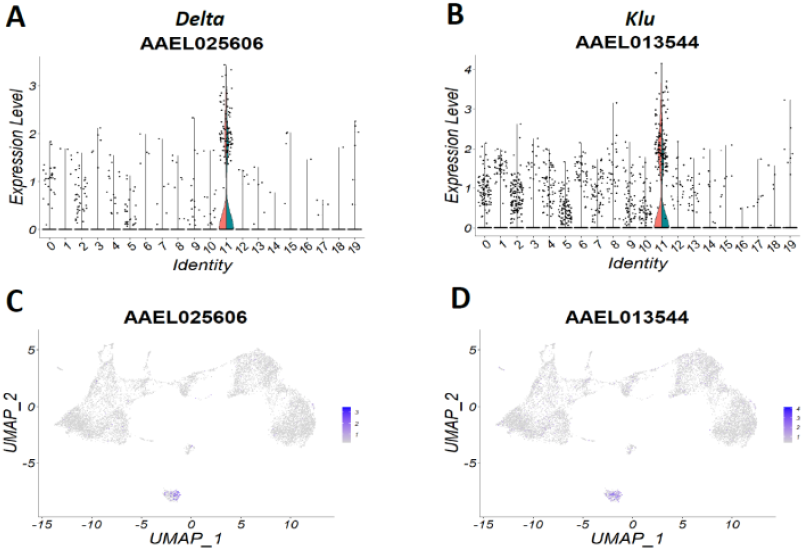
Identification of the intestinal stem cell/enteroblast (ISC/EB) cluster in the female midgut of *Ae. aegypti*. (**A**) VlnPlot showing *Delta* expression levels. Data points within or near the red zone represent single nuclei from the blood-fed midguts, those within or near the green zone are single nuclei from the sugar-fed midguts. (**B**) VlnPlot showing *Klu* expression levels. Data points within or near the red zone represent single nuclei from the blood-fed midguts, those within or near the green zone are single nuclei from the sugar-fed midguts. (**C**) FeaturePlot showing *Delta* expression levels. (**D**) FeaturePlot showing *Klu* expression levels. Shown expression levels are values based on LnFoldChange. “Identity” represents the individual cell clusters.

**Fig. S3.**
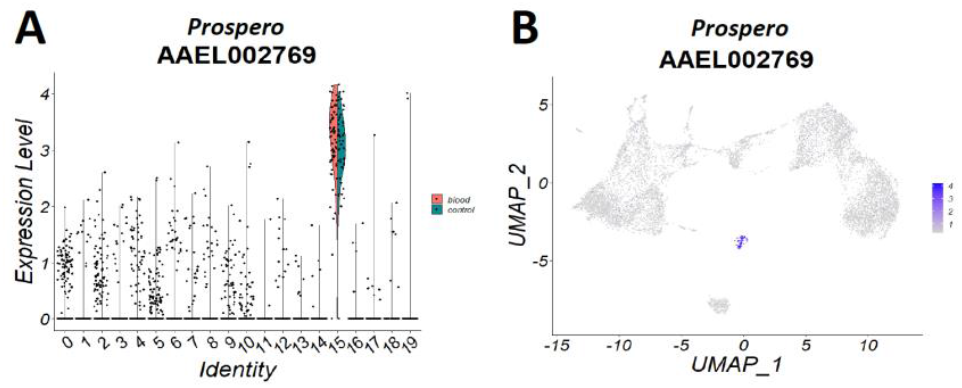
Identification of the enteroendocrine cell cluster (EE) in the female midgut of *Ae. aegypti*. (**A**) VlnPlot showing *Prospero* expression levels. Data points within or near the red zone represent single nuclei from the blood-fed midguts, those within or near the green zone are single nuclei from the sugar-fed midguts. (**B**) FeaturePlot showing *Prospero* expression levels. Shown expression levels are values based on LnFoldChange. “Identity” represents the individual cell clusters.

**Fig. S4.**
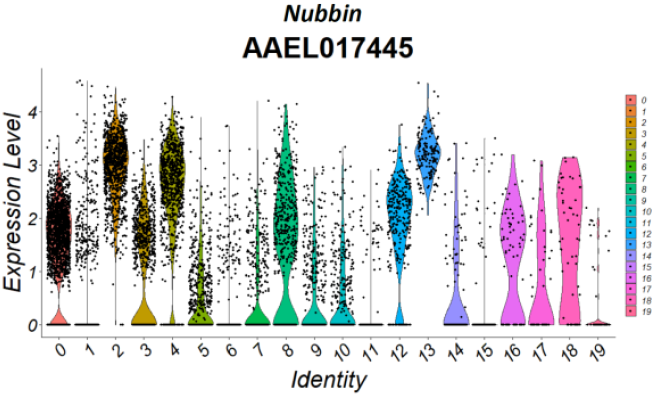
Identification of the enterocyte (EC) cell clusters in the female midgut of *Ae. aegypti*. VlnPlot showing expression levels (average LnFoldChange) of *Nubbin*. “Identity” represents the individual cell clusters.

**Fig. S5.**
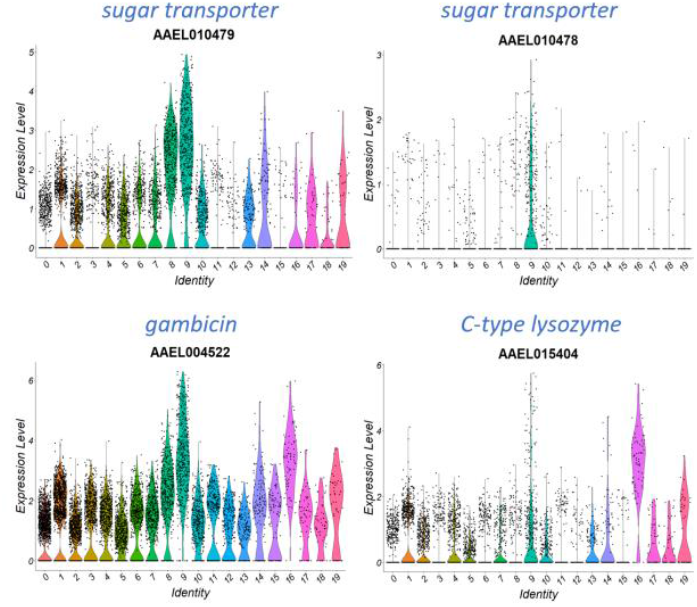
Identification of cardia cell clusters in the female midgut of *Ae. aegypti*. VlnPlots showing expression levels of two sugar transporters, *gambicin, and C-type lysozyme*. Shown expression levels are values based on LnFoldChange. “Identity” represents the individual cell clusters.

**Fig. S6.**
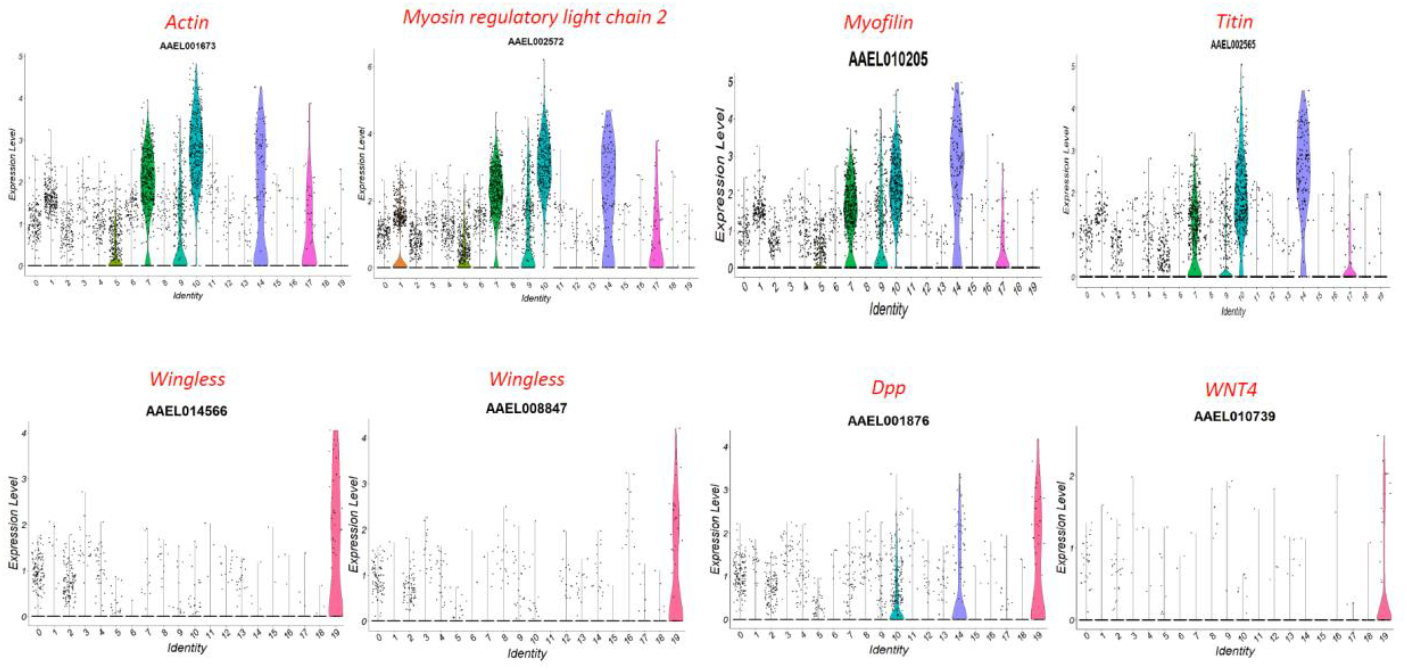
Identification of visceral muscle cell clusters (VM) in the female midgut of *Ae. aegypti*. VlnPlots showing the expression levels of actin, *myosin regulatory light chain 2, myofilin, titin, Wingless, Dpp*, and *WNT4*. Shown expression levels are values based on LnFoldChange. “Identity” represents the individual cell clusters.

**Fig. S7.**
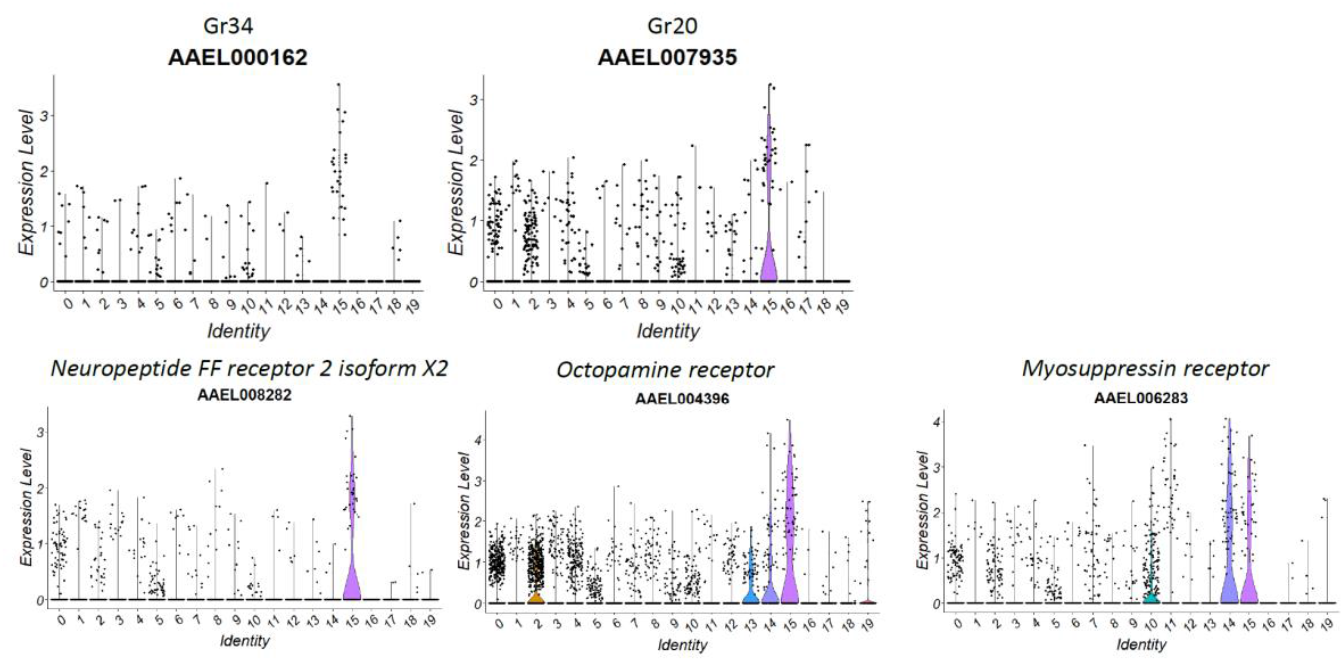
VlnPlots showing the expression levels of the two gustatory receptors (Gr34 and Gr20), *Neuropeptide FF receptor 2 isoform X2, Octopamine receptor*, and *Myosuppressin receptor* in different midgut cell clusters. Shown expression levels are values based on LnFoldChange. “Identity” represents the individual cell clusters.

**Fig. S8.**
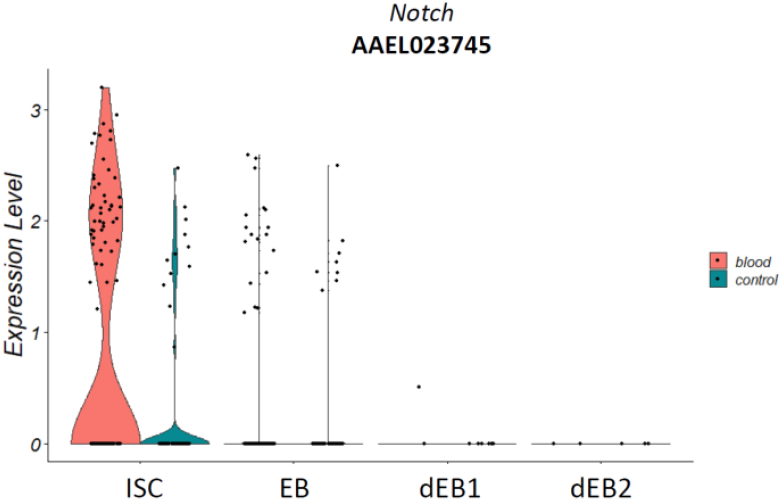
VlnPlot showing the changes in *Notch* expression levels in ISC, EB, dEB-1, and dEB-2 as a consequence of blood meal ingestion. Shown expression levels are values based on LnFoldChange.

